# Redox potential as a master variable controlling pathways of metal reduction by *Geobacter sulfurreducens*

**DOI:** 10.1101/043059

**Authors:** Caleb E. Levar, Colleen L. Hoffman, Aubrey J. Dunshee, Brandy M. Toner, Daniel R. Bond

## Abstract

*Geobacter sulfurreducens* uses at least two different pathways to transport electrons out of the inner membrane quinone pool before reducing acceptors beyond the outer membrane. When growing on electrodes poised at oxidizing potentials, the CbcL-dependent pathway operates at or below redox potentials of −0.10 V vs. the Standard Hydrogen Electrode (SHE), while the ImcH-dependent pathway operates only above this value. Here, we provide evidence that *G. sulfurreducens* also requires different electron transfer proteins for reduction of a wide range of Fe(III)- and Mn(IV)- (oxyhydr)oxides, and must transition from a high- to low-potential pathway during reduction of commonly studied soluble and insoluble metal electron acceptors. Freshly precipitated Fe(III)-(oxyhydr)oxides could not be reduced by mutants lacking the high potential pathway. Aging these minerals by autoclaving did not change their powder X-ray diffraction pattern, but restored reduction by mutants lacking the high-potential pathway. Mutants lacking the low-potential, CbcL-dependent pathway had higher growth yields with both soluble and insoluble Fe(III). Together, these data suggest that the ImcH-dependent pathway exists to harvest additional energy when conditions permit, and CbcL switches on to allow respiration closer to thermodynamic equilibrium conditions. With evidence of multiple pathways within a single organism, the study of extracellular respiration should consider not only the crystal structure or solubility of a mineral electron acceptor, but rather the redox potential, as this variable determines the energetic reward affecting reduction rates, extents, and final microbial growth yields in the environment.

## Introduction

Fe(III)-(oxyhydr)oxides can exist in at least 15 mineral forms with variable physiochemical properties that span a wide range of formal oxidation-reduction (redox) midpoint potentials (Majzlan, Navrotsky, & Schwertmann, 2004; Majzlan, 2011, 2012; Nealson & Saffarini, 1994; Schwertmann & Cornell, 2000; Thamdrup, 2000). The electron-accepting potential of any given Fe(III)-(oxyhydr)oxide structure is not a fixed value, and becomes less favorable with increasing crystallinity, particle size, pH, or ambient Fe(II) concentration (Majzlan, 2012; Navrotsky, Mazeina, & Majzlan, 2008; Sander, Hofstetter, & Gorski, 2015; Thamdrup, 2000). Differences in effective redox potential alters the energy available to be captured by bacteria able to couple the oxidation of an electron donor to reduction of these minerals (Flynn, O’Loughlin, Mishra, DiChristina, & Kemner, 2014; Thauer, Jungermann, & Decker, 1977). Because of this structural and environmental diversity, organisms able to reduce metals could sense the energy available in their electron acceptors and utilize different electron transfer pathways, similar to how *E. coli* uses distinct terminal oxidases in response to levels of available oxygen (Green & Paget, 2004; Russell & Cook, 1995).

One well-studied dissimilatory metal reducing organism, *Geobacter sulfurreducens,* can grow via reduction of metal (oxyhydr)oxides ranging in predicted midpoint redox potential from +0.35 V vs. the Standard Hydrogen Electrode (SHE)(e.g., birnessite, ca. Na_x_Mn_2-x_(IV)Mn(III)_x_O_4,_ x ~ 0.4) to −0.17 V vs. SHE (e.g., goethite, α-FeOOH) (Caccavo et al., 1994; Majzlan et al., 2004; Majzlan, 2012; Post & Veblen, 1990; Thamdrup, 2000). Recent work examining electron transfer from *G. sulfurreducens* to poised graphite electrodes demonstrated that this organism uses at least two different inner membrane electron transfer pathways, known as the CbcL- and ImcH-dependent pathways (Levar, et al., 2014; Zacharoff, et al., 2016). The ‘low’ potential, CbcL-dependent pathway, is required for growth with electrodes at or below potentials of −0.10 V vs. SHE, while the ‘high’ potential, ImcH-dependent pathway, is essential when electrodes are poised at redox potentials above this value. As mutants lacking each pathway only grow with electrodes poised above or below these thresholds, cells containing only CbcL or ImcH might also be able to be used as ‘sensors’ to report the redox potential of an extracellular electron acceptor such as a metal (oxyhydr)oxide. As it is difficult to determine the effective redox potential of these minerals (Sander et al., 2015), such information could aid laboratory characterization, and provide evidence that bacteria use different pathways for different metals in the environment.

The discovery of multiple inner membrane pathways is based on work with electrodes held at constant potentials, but poses many questions regarding *G. sulfurreducens’* interactions with more complex mineral substrates. Does the organism transition from one pathway to the other as simple environmental factors such as Fe(II):Fe(III) ratios alter redox potentials? Do manipulations known to alter redox potential, such as pH or particle size, also influence the pathway utilized? As this +/- 0.5 V redox potential range represents nearly 50 kJ per mol of electrons transferred, do different electron transfer mechanisms allow higher yield of this microbe?

Here, we demonstrate that *G. sulfurreducens* requires both the CbcL- and the ImcH-dependent electron transfer pathways for complete reduction of a variety of Fe and Mn minerals. By using mutants only able to function above or below specific redox potentials, we show that minerals often used for study of extracellular electron transfer begin as ‘high’ potential electron acceptors reducible by the ImcH-dependent pathway, but transition during reduction to ‘low’ potential electron acceptors requiring the CbcL-dependent pathway. Simple variations in mineral handling such as autoclaving, or pH changes that alter redox potential by 30 mV, can alter the electron transfer pathway required by the organism, further showing that bacteria respond to subtle changes in mineral redox potentials. These data highlight the complexity of studying growth with minerals, and suggests that the proteins used for electron flow out of *G. sulfurreducens* are more strongly influenced by redox potential than the crystal structure, aggregation behavior, or even solubility of the terminal acceptor.

## Materials and methods

### Bacterial strains and culture conditions

Strains of *Geobacter sulfurreducens* used in this study are described in Table 1. All strains were routinely cultured from freezer stocks stored at −80 degrees Celsius, separate colony picks from agar plates were used to initiate replicate cultures used in all experiments. Strains were cultured in NB minimal medium composed of 0.38 g/L KCl, 0.2 g/L NH_4_Cl, 0.069 g/L NaH_2_PO_4_∙H_2_O, 0.04 g/L CaCl_2_∙2H_2_O, and 0.2 g/L MgSO_4_∙7H_2_O. 10 ml/L of a chelated mineral mix (chelator only used for routine growth with soluble acceptors, see below) containing 1.5g/L NTA, 0.1 g/L MnCl_2_∙4H_2_O, 0.3 g/L FeSO_4_∙7H_2_O, 0.17 g/L CoCl_2_∙6H2O, 0.1 g/L ZnCl_2_, 0.04 g/L CuSO_4_∙5H2O, 0.005 g/L AlK(SO_4_)_2_∙12H_2_O, 0.005g/L H_3_BO_3_, 0.09 g/L Na_2_MoO_4_, 0.12 g/L NiCl_2_, 0.02 g/L NaWO_4_∙2H2O, and 0.1 g/L Na_2_SeO_4_ was also added. Fumarate (40 mM) was used as an electron acceptor for growth of initial stocks picked from colonies, and acetate (20 mM) was used as the sole electron and carbon source. Unless otherwise noted, the pH of the medium was adjusted to 6.8 and buffered with 2 g/L NaHCO_3_, purged with N_2_:CO_2_ gas (80%/20%) passed over a heated copper column to remove trace oxygen, and autoclaved for 20 minutes at 121 degrees Celsius.

**Table 1.**
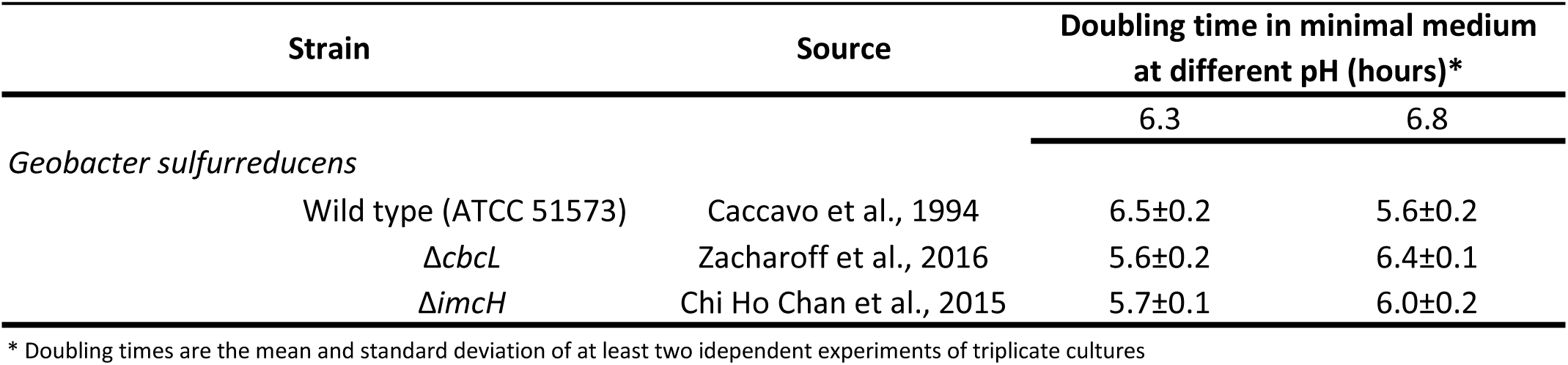

### Growth of *G. sulfurreducens* with Fe(III)-citrate and continuous redox potential monitoring

Minimal medium containing 20 mM acetate as the carbon and electron donor and ~80 mM Fe(III)-citrate as the electron acceptor was added to sterile bioreactors constructed as previously described, using both working and counter electrodes bare platinum wire approximately 2 cm in length. The headspace of each reactor was purged with anaerobic and humidified N_2_:CO_2_ (80%/20%) gas. Calomel reference electrodes were used, and a solution of 0.1 M NaSO_4_ stabilized with 1% agarose separated from the medium by a vycor frit provided a salt bridge between the reference electrode and the growth medium. The working volume of the bioreactors was 15 ml. A 16-channel potentiostat (VMP; Bio-Logic, Knoxville, TN) using the EC-lab software (v9.41) was used to measure the open circuit voltage between the working and reference electrode every 500 seconds, providing a continuous readout of the redox potential in the medium. After redox potential was stable for at least 5 hours, stationary phase cells were added to each bioreactor at a 1:100 v/v inoculum size and the change in redox potential was measured over time. When co-cultures were used, and equal volume of cells was added for each strain (i.e., 1:100 each of Δ*cbcL* and Δ*imcH* was added after the voltage was stable).

#### X-ray diffraction (XRD) measurements

Approximately 0.2-0.5 g of untreated bulk mineral sample or 2.5 ml of mineral suspension in medium were analyzed by powder XRD. Minerals in basal media were separated from suspension by filtration (0.22 μm) in an anaerobic chamber (Coy Laboratory Products) under a N_2_:CO_2_:H_2_ (75%:20%:5%) atmosphere, and stored in sealed mylar bags at −20 degrees Celsius prior to analysis. Diffractograms were measured from using 20° to 89° (2Θ) range, a step size of 0.02 2Θ, and a dwell of 0.5°/minute using a Rigaku MiniFlex ASC-6AM with a Co source. The resulting diffractograms were background subtracted, and phases were identified using Jade v9.1 software (MDI, Inc.).

#### Preparation of Fe(III)-(oxyhydr)oxides

“Poorly crystalline FeO(OH)” was produced after Lovley and Phillips, 1986, by adding 25% NaOH dropwise over 90 minutes to a rapidly stirring 0.4 M solution of FeCl_3_ until the pH was 7.0. The solution was held at pH 7.0 for one hour, and the resulting suspension was washed with one volume of de-ionized and distilled water and used immediately (<24 after synthesis) for XRD analysis and growth studies as rapid aging of such products to more crystalline minerals (e.g., goethite) has been observed even when kept in the dark at 4 degrees Celsius (Raven, Jain, & Loeppert, 1998). The resulting mineral prior to its addition to medium and autoclaving (“untreated mineral”) was identified as akaganeite (β-FeOOH) by powder X-ray diffraction. Washing this “untreated mineral” samples with up to three additional volumes of de-ionized and distilled water did not change the XRD pattern of the mineral or the reduction phenotypes observed. For all minerals, “untreated” refers to the mineral suspension in MiliQ water prior to addition to medium and autoclaving, “fresh” refers to the mineral suspension in medium, and “autoclaved” samples result from adding the “untreated” minerals to medium and autoclaving (ie, autoclaving the “fresh” samples).

Goethite and 2-line ferrihydrite (ca. Fe_10_O_14_(OH)_2_) (Michel et al., 2007) were synthesized after Schwertmann and Cornell, 2000, with some modifications. For goethite, a solution of FeCl_3_ was precipitated through neutralization with 5 N KOH, and the resulting alkaline mineral suspension was aged at 70 degrees Celsius for 60 hours to facilitate goethite formation. The suspension was then centrifuged, decanted, and washed with de-ionized and distilled water prior to freeze drying. For 2-line ferrihydrite, a solution of FeCl_3_ was neutralized with KOH until the pH was 7.0. Suspensions were rinsed with de-ionized and distilled water and either freeze dried or centrifuged and suspended in a small volume of de-ionized and distilled water to concentrate the final product. Freeze dried samples yielded XRD patterns indicative of 2-line ferrihydrite, but suspension of hydrated samples in growth medium had an altered XRD pattern more similar to that of akaganeite. Freeze dried samples were stored at −80 degrees Celsius prior to addition to growth medium, while hydrated samples were used within 24 hours of synthesis.

Schwertmannite (Fe_8_O_8_(OH)_6_(SO_4_)·nH_2_O) was synthesized using a rapid precipitation method involving the addition of a 30% solution of H_2_O_2_ to a solution of FeSO_4_ (Regenspurg, Brand, & Peiffer, 2004). The mixture was stirred rapidly for at least 12 hours until the pH was stable, at ~2.35. The solids were allowed to settle for 1 hour, and the supernatant carefully decanted. Solids were washed by centrifuging at 3,700 x g and suspending in 1 volume of de-ionized and distilled water. The product was concentrated by a final centrifugation at 3,700 x g for 5 minutes and resuspension in 1/20 the initial volume. XRD identified this initial product as schwertmannite, though subsequent addition to media at neutral pH values and autoclaving rapidly altered the mineral form.

#### Iron reduction assays

Basal media were prepared as above with some modifications. Fumarate was omitted as the Fe(III)-(oxyhydr)oxides were desired as the sole electron acceptor. NTA free trace mineral mix (in which all components were first dissolved in a small volume of HCl) was used in order to eliminate exogenous chelating compounds from the media. All media were purged with N_2_:CO_2_ gas (80%:20%) passed over a heated copper column to remove trace oxygen. Where indicated, media were autoclaved for 20 minutes at 121 degrees Celsius on a gravity cycle, and were immediately removed to cool at room temperature in the dark. As is standard for *G. sulfurreducens* medium containing Fe(III)-(oxyhydr)oxide (Caccavo et al., 1994), additional NaH_2_PO_4_∙H_2_O (to a final concentration of 0.69 g/L) was added prior to autoclaving, as more recalcitrant and less reducible forms of Fe(III)-(oxyhydr)oxide will form when autoclaved without additional phosphate. For fresh akaganeite and 2-line ferrihydrite, 1 ml of the Fe-(oxyhydr)oxide suspension was added to 9 ml of basal medium. The final pH was adjusted by altering the pH of the basal medium prior to mineral addition and by altering the concentration of NaHCO_3_ used prior to purging with anaerobic N_2_:CO_2_ gas (80%/20%). For goethite, an 88 g/L suspension was made in de-ionized and distilled water and 1 ml of this suspension was added to 9 ml of basal medium. The resulting medium had an effective Fe_total_ concentration of ~20 mM as determined by fully reducing an acidified sample with hydroxylamine and measuring the resulting Fe(II) using a modified Ferrozine assay (Lovley & Phillips, 1987). Because the synthesis of schwertmannite results in an acidic product below the pH at which *G. sulfurreducens* grows optimally (Caccavo et al., 1994; Regenspurg et al., 2004), the mineral suspension was added to the basal medium (1:9) and the pH of the solution was adjusted with NaOH and bicarbonate to produce a final pH of 6.8. For freeze dried 2-line ferrihydrite, 0.2 grams of freeze dried sample was added per 10 ml of basal medium.

In all cases, 100 μl of electron acceptor-limited stationary phase cells (OD_600_ = 0.55) grown from single colony picks in NB basal medium containing acetate (20 mM) and fumarate (40 mM) were used to inoculate the Fe(III)-(oxyhydr)oxide containing media, with acetate provided as the electron and carbon donor and the Fe(III)-(oxyhydr)oxide as the sole electron acceptor.

For all Fe(III)-(oxyhydr)oxide incubations except those involving goethite, a small sample was removed at regular intervals and dissolved in 0.5 N HCl for at least 24 hours at 23-25 degrees Celsius in the dark before the acid extractable Fe(II) present in the sample was measured. For incubations with goethite, samples were incubated in 1 N HCl at 65 degrees Celsius for at least 24 hours in the dark, followed by centrifugation at 13,000 x g for 10 minutes to remove solids. Fe(II) in the supernatant was then measured.

#### Manganese reduction assays

Birnessite (Na_x_Mn_2-x_(IV)Mn(III)_x_O_4,_ x ~ 0.4) was prepared using the protocol described in Chan, et al, 2015. Briefly, a solution of KMnO_4_ was added to an equal volume of MnCl_2_·4H_2_O solution and mixed vigorously. Solids were allowed to settle, and the overlaying liquid was decanted off. The resulting solid was washed in de-ionized and distilled water and concentrated by centrifugation. This suspension was added (~20mM) to basal medium containing acetate (10 mM) as the carbon and electron donor. Cells were inoculated to a calculated OD_600_ of 0.005 from electron acceptor-limited stationary phase cells. Mn(II) was measured indirectly as previously described (Chan et al., 2015; Levar et al., 2014). Briefly, samples were acidified in 2 N HCl with 4 mM FeSO_4_ added and allowed to dissolve overnight at 25 degrees Celsius in the dark. After all solids were dissolved, the Fe(II) concentration was measured. Because the reduction of Mn(IV) by Fe(II) is thermodynamically favorable, the measured Fe(II) concentration can be used to extrapolate the Mn(II) present a given sample. When measured Fe(II) concentrations are high, this implies that Mn(IV) concentrations are low (ie, Mn(IV) has been reduced by the bacterial culture).

## Results

The inner membrane multiheme cytochromes ImcH and CbcL are implicated in transfer of electrons out of the quinone pool when electron acceptors fall within different redox potential windows (Levar et al., 2014; Zacharoff et al., 2016), though it is important to note that direct roles for ImcH and CbcL in the movement of electrons out of the quinone pool have not been demonstrated. Mutants lacking *imcH* are only able to reduce poised electrodes with ‘low’ redox potentials (≤ −0.10 V vs. SHE), and are deficient in reduction of electrodes at redox potentials above this value (Levar et al., 2014). In contrast, *cbcL* mutants have the opposite phenotype, reducing electrodes with relatively high electrode redox potentials, and having a deficiency in reduction of electrodes at or below −0.10 V vs. SHE (Zacharoff et al., 2016). The genes for ImcH and CbcL are constitutively expressed (Zacharoff et al., 2016), yet use of each pathway appears to switch on or off in a matter of minutes during sweeps of increasing or decreasing potential known as catalytic cyclic voltammetry (Marsili, Rollefson, Baron, Hozalski, & Bond, 2008; Yoho, Popat, & Torres, 2014; Zacharoff et al., 2016). Genetic complementation of each pathway has been demonstrated using a variety of extracellular electron acceptors (Chan et al., 2015; Levar et al., 2014; Zacharoff et al., 2016).

Because it is difficult to measure the effective redox potential of metal (oxyhydr)oxides (Sander et al., 2015), the redox potential of cultures undergoing active reduction of soluble Fe(III)-citrate was first monitored for each strain (Figure 1). While the wild type reduced Fe(III) completely to Fe(II), achieving a redox potential of −0.255 V vs. SHE, the Δ*cbcL* mutant strain (lacking the low-potential system) was unable to reduce all Fe(III), and the redox potential only lowered to −0.15 V. The Δ*imcH* mutant strain (lacking the high potential system) was unable to reduce Fe(III) when inoculated into the high-potential environment. When both Δ*cbcL* and Δ*imcH* mutant strains were inoculated into the same reactor, redox potential decreased with similar kinetics as the Δ*cbcL* strain, but continued on to a final value similar to wild type. This was consistent with the Δ*cbcL* mutant strain first reducing Fe(III), until the redox potential was low enough to be usable by the Δ*imcH* mutant. This hard transition from where the Δ*cbcL* and Δ*imcH* mutant strains could actively reduce Fe(III) occurred in a range (between 0 and −0.1 V) was similar to the theoretical potentials of many Fe(III)-(oxyhydr)oxides. With that observation that these mutants behaved with Fe(III) citrate in a manner comparable to electrodes, we hypothesized that mutants lacking the ImcH- and CbcL-dependent electron transfer pathways would respond similarly when exposed to commonly used metal (oxyhydr)oxides.

**Figure 1.**
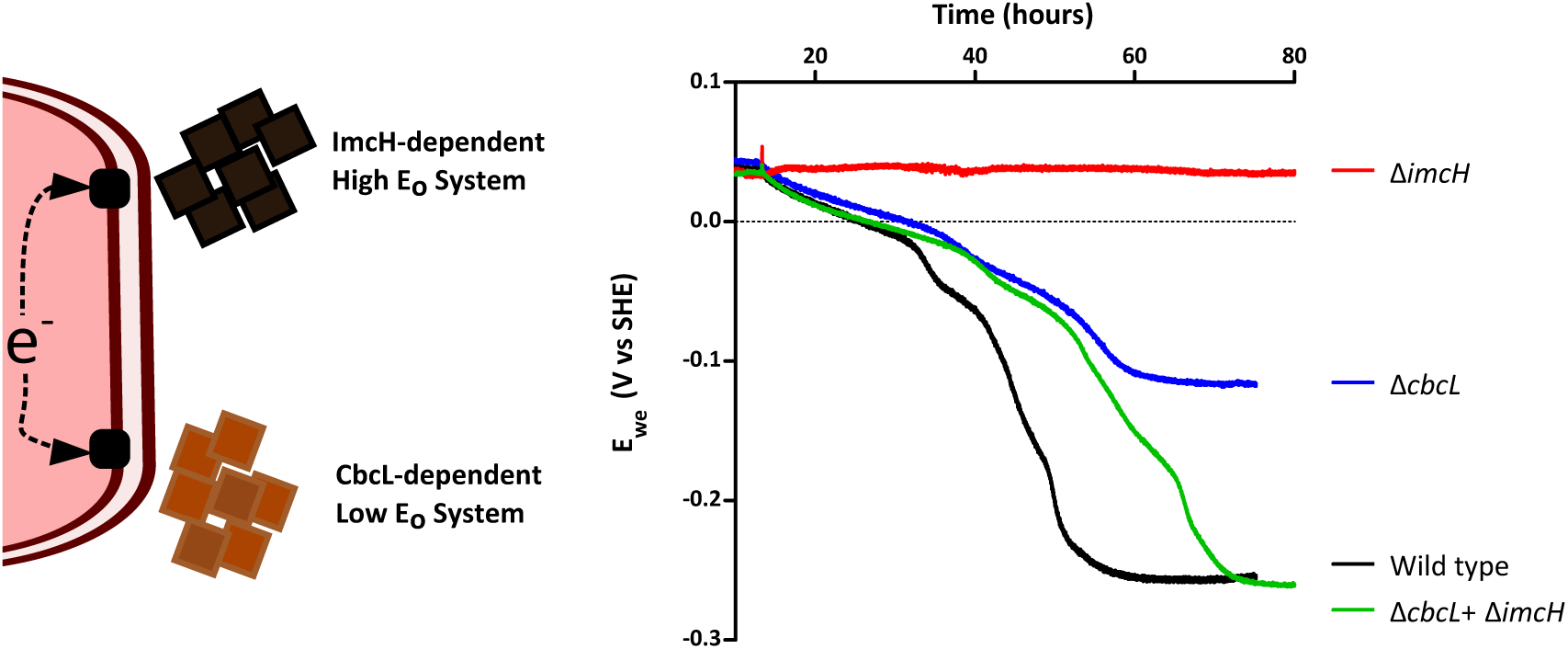
*Redox potential controls the behavior of Δ*imcH *and Δ*cbcL *mutants*. Redox potential was recorded in real time during reduction of Fe(III)-citrate by each culture. Cells lacking the CbcL-dependent pathway (Blue trace) initially reduced Fe(III), but could not lower the redox potential below −0.15 V vs SHE. Cells lacking the ImcH-dependent pathway (Red trace) could not lower the redox potential from the uninoculated value (+0.04 V vs SHE). The redox potential was lowed to the same point as wild type (black trace) when a co-inoculum of the two mutants was used (Green trace). Traces are representative of at least triplicate experiments for each strain or co-culture.

### Evidence that the electron transfer pathway changes even in the absence of XRD-pattern changes

Most Fe(III)-(oxyhydr)oxides do not exist at the high or low extremes represented by electrodes or fresh Fe(III) citrate, but possess electron accepting potentials predicted to lie near the −0.10 V redox potential threshold that requires a transition from ImcH-dependent to the CbcL-dependent reduction pathways in electrode- and Fe(III)-citrate grown cells cells (Levar et al., 2014; Majzlan et al., 2004; Majzlan, 2012; Thamdrup, 2000; Zacharoff et al., 2016). While testing this with electrodes or soluble electron acceptors is straightforward, the redox potential of Fe(III)-(oxyhydr)oxides in this range is not easy to predict or measure (Sander et al., 2015). Aging (accelerated by autoclaving or freeze drying) can increase particle size and lower redox potential, or drive re-crystallization to lower redox potential forms (Navrotsky et al., 2008; Roden, Urrutia, & Mann, 2000; Roden & Zachara, 1996; Roden, 2006; Thamdrup, 2000). As Fe(III)-(oxyhydr)oxide reduction is a proton consuming reaction (Kostka & Nealson, 1995; Majzlan, 2012; Thamdrup, 2000), for each unit decrease in pH, the redox potential of the Fe(III)/Fe(II) redox couple increases by ~59 mV per the Nernst equation (Bonneville, Van Cappellen, & Behrends, 2004; Kostka & Nealson, 1995; Majzlan, 2012; Thamdrup, 2000).

To test the hypothesis that *G. sulfurreducens* also utilizes ImcH- vs. CbcL-dependent pathways to reduce minerals on the basis of redox potential, we prepared Fe(III)-(oxyhydr)oxides by slow precipitation of FeCl_3_ with NaOH (Lovley & Phillips, 1986), and characterized minerals before and after addition to medium using powder XRD. While Fe(III)-(oxyhydr)oxides prepared in this manner are often referred to as “poorly crystalline” or “amorphous”, all XRD signatures of our slow-precipitation preparations were consistent with akaganite (β-FeOOH) (Figure 2E). Because the β-FeOOH/Fe(II) redox couple has a formal midpoint potential near −0.10 V vs. SHE (Majzlan, 2012), a fully oxidized suspension of akaganeite would initially have a redox potential well above 0 V vs SHE, and theoretically require only the ‘high’ potential pathway of *G. sulfurreducens.*

**Figure 2.**
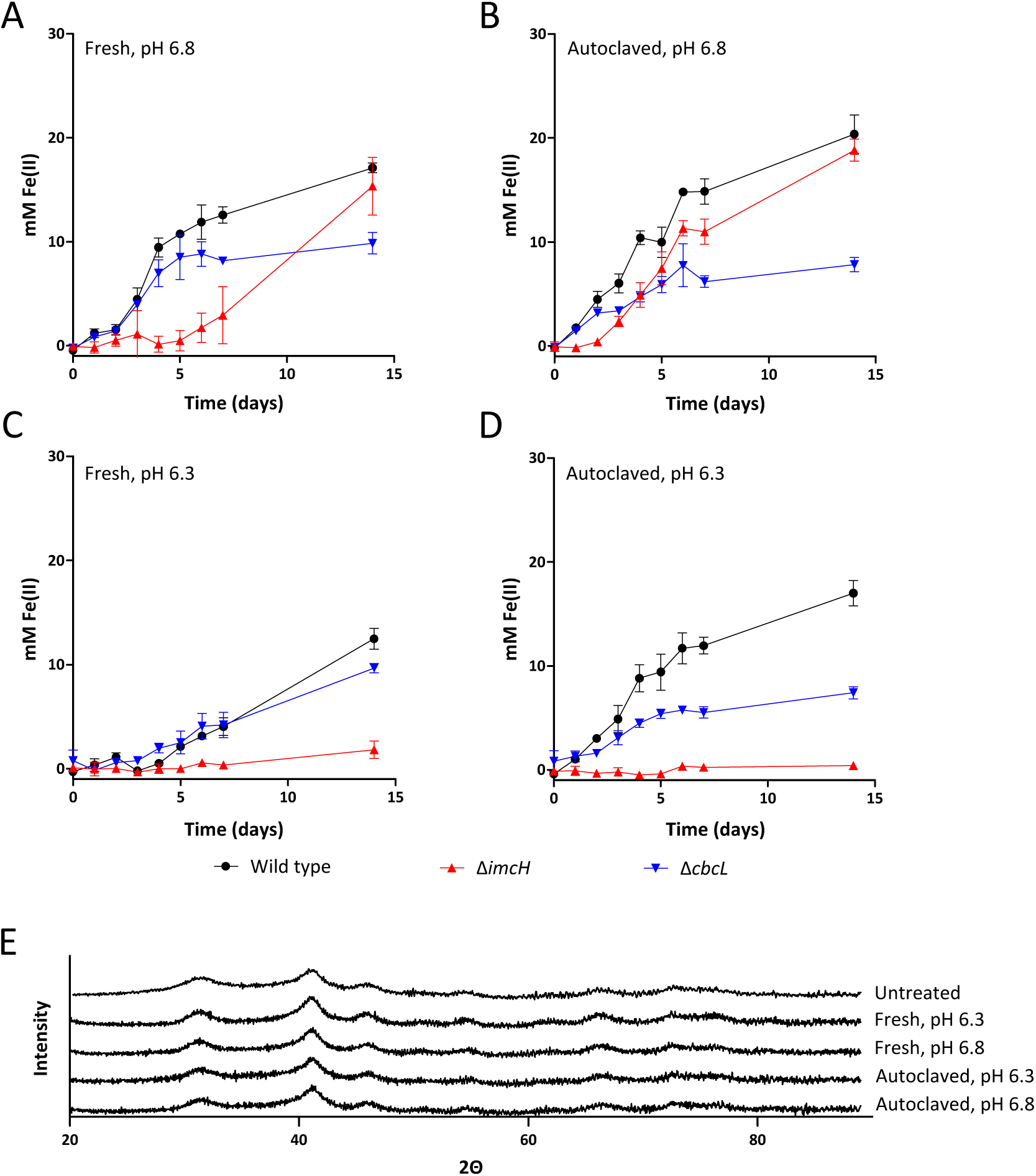
*Autoclaving and changing pH alters mutant phenotypes*. In all cases, wild type is represented by black circles, Δ*cbcL* by downward pointing blue triangles, and Δ*imcH* by upward pointing red triangles. (A) A long lag is observed for Δ*imcH* inoculated into medium with no autoclaving (“Fresh”). (B) The lag observed for Δ*imcH* is decreased when the medium is aged through autoclaving. (C) Decreasing the pH by 0.5 units eliminates akaganeite reduction by Δ*imcH*. (D). Aging pH 6.3 medium through autoclaving is not sufficient to decrease the redox potential enough to allow for reduction by Δ*imcH*. All data shown are the mean and standard deviation for triplicate cultures, and are representative of at least duplicate incubations. (E) XRD patterns of the mineral used in panels A-D. The XRD pattern for the mineral prior to addition to medium and autoclaving (“Untreated”) is also shown.

When wild-type, Δ*imcH,* and Δ*cbcL* strains were incubated with akaganeite that had not been autoclaved (labeled as “Fresh”), wild-type cultures reduced the provided Fe(III)-(oxyhydr)oxide over the course of 15 days. The Δ*cbcL* cultures containing the high-potential pathway initially reduced fresh akaganeite at rates similar to wild type, but reduction slowed as Fe(II) accumulated, and the concentration of Fe(II) never reached the extent of wild type. In contrast, Δ*imcH* cultures lacking the high potential pathway demonstrated a substantial lag, remaining near background for 7 days (Figure 2A). Once metal reduction began, Δ*imcH* cultures were able to reach the final extent observed in wild type. 14 day cell free controls did not exhibit any observable Fe(III)-(oxyhrdr)oxide reduction.

Medium containing akaganeite was then autoclaved, a treatment predicted to lower redox potential due to accelerated aging. Both wild type and Δ*cbcL* performed similar to the fresh, unautoclaved sample, but the lag observed with Δ*imcH* was now much shorter, consistent with a lower redox potential (Figure 2B). Despite the change in the Δ*imcH* Fe(III)-(oxyhydr)oxide reduction phenotype, the XRD pattern before and after autoclaving was similar (Figure 2E). Such a lack of XRD alteration in response to autoclaving has been previously noted (Hansel et al., 2015). This suggests that phenotypic changes were due to mineral alteration outcomes not detected by powder XRD, such as short-range structural order or particle aggregation, that lowered effective redox potential and allowed growth of the Δ*imcH* strain lacking the high-potential pathway.

### Reduction phenotypes also respond to changes in medium pH that alter redox potential

The lag in extracellular electron transfer by Δ*imcH,* in contrast to immediate reduction by Δ*cbcL*, suggests that fresh akaganeite has an initial redox potential greater than −0.10 V vs. SHE. Based on this hypothesis, after many days, a small amount of Fe(II) accumulates, and potential drops into a range reducible by Δ*imcH.* If aging fresh akaganite by autoclaving shortened the lag in Figure 2B by lowering redox potential, then manipulations designed to raise redox potential should extend this lag or prevent Δ*imcH* from reducing akaganeite altogether.

When the pH of basal medium containing akaganeite was adjusted to 6.3 (raising the redox potential ~30 mV (Kostka & Nealson, 1995; Majzlan, 2012; Thamdrup, 2000)), near complete inhibition of reduction by Δ*imcH* was observed (Figure 2C). Autoclaving pH 6.3 akaganeite medium was not sufficient to restore the reduction phenotype of Δ*imcH,* but it did aid reduction by Δ*cbcL* (Figure 2D). As before, autoclaving at 6.3 did not substantially alter the XRD pattern of the mineral provided (Figure 2E). Importantly, wild type, Δ*imcH,* and Δ*cbcL* grew without significant defects at pH values of 6.3 or 6.8 in control experiments containing fumarate as the sole terminal electron acceptor (Table 1). Based on these results, when akaganeite is autoclaved, the initial redox potential is likely lowered closer to −0.10 V vs. SHE, permitting operation of the CbcL-dependent, low-potential pathway. Fresh akaganeite, especially at pH 6.3, behaves consistent with a redox potential of 0 V or greater, thus preventing reduction by cells containing only the low-potential pathway.

These two experiments, using aging to reduce redox potential and pH to raise redox potential, caused changes in Δ*imcH* phenotypes that provided new evidence that *G. sulfurreducens* discriminates between extracellular electron acceptors independent of the mineral properties accessed by powder XRD. They also pointed to best practices for medium preparation in order to obtain consistent results if the phenotype is sensitive to the redox potential of the environment. The amount of Fe(II) accumulated after 7 days provided a repeatable readout of the ability of a *G. sulfurreducens* strain or mutant reduce a given Fe(III)-(oxyhydr)oxide. According to this assay, the more Fe(II) produced in 7 days by an Δ*imcH* mutant, the lower the initial redox potential of the mineral likely was.

### Other mineral forms consistent with a multiple-pathway model

Additional Fe(III)-(oxyhydr)oxide minerals were then synthesized to test interactions of these variables. When schwertmannite was synthesized using a rapid precipitation method (Regenspurg et al., 2004), the mineral was stable at pH values below pH 4.0 and its structure could be confirmed by XRD. However, abiotic transformation of schwertmannite to other mineral forms is accelerated at pH values greater than 4.5 (Blodau & Knorr, 2006) and analysis of XRD patterns after schwertmannite was added to *Geobacter* growth medium revealed rapid formation of XRD-amorphous minerals (Figure 3E), though structural information may be elucidated with more powerful techniques such as synchrotron EXAFS. Reduction phenotypes suggested this fresh mineral also produced a redox potential near 0 V vs. SHE, as cells lacking the high potential pathway (Δ*imcH*) showed a lag at pH 6.8, which was shorter after the mineral was autoclaved (Figure 3B). Reduction by Δ*imcH* was worst when the redox potential was further raised by buffering to pH 6.3 (Figure 3C), while autoclaving pH 6.3 medium allowed a small amount of Fe(II) accumulation that ended the lag after >8 days. These patterns with schwertmannite-based minerals were similar to those observed with akaganite.

**Figure 3.**
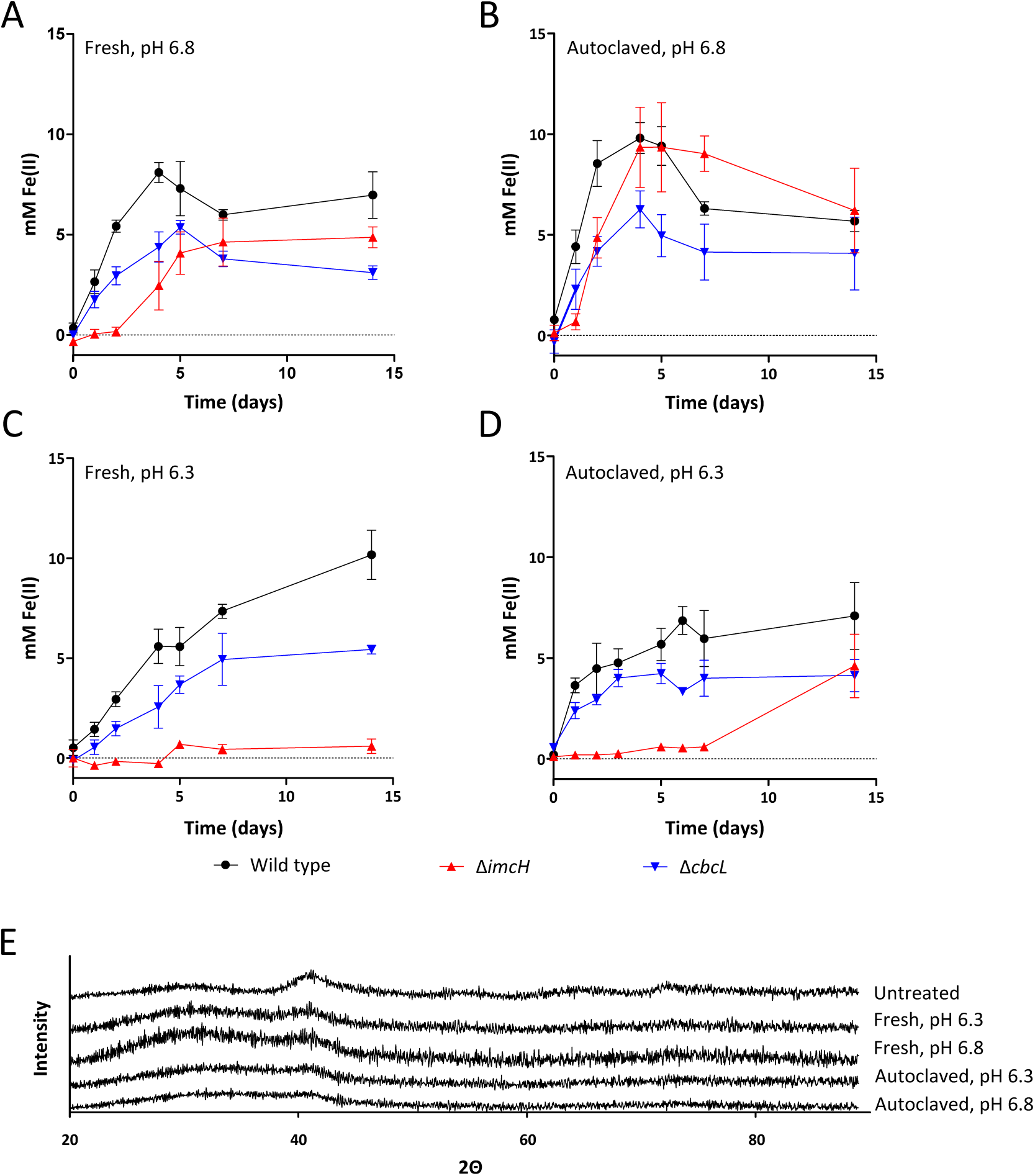
*Addition of schwertmannite to growth medium at pH 6.3 or pH 6.8 alters mineralogy, and mutant behavior*. Rapid precipitation of an Fe sulfide solution with hydrogen peroxide yields XRD-pure schwertmannite (“untreated”, Panel E), but when this mineral suspension is added to growth medium, abiotic transformation is observed. Autoclaving these media lead to further mineral transformation. Treatments aimed at raising and lowering redox potential affected reduction by *ΔimcH* mutants similar to results seen in Figure 2.

When Fe(III)-(oxyhrdr)oxide was prepared according to the 2-line ferrihydrite protocol of Schwertmann and Cornell (2000), wild type and mutant phenotypes were again what was observed with akaganite and schwertmannite. Freshly prepared mineral was reduced by Δ*cbcL* to within 67% of wild type after seven days. When this mineral was aged by freeze drying (resulting in XRD-confirmed 2-line ferrihydrite), reduction by Δ*cbcL* only 30% of wild type, suggesting that freeze drying lowered redox potential of the mineral to a point where the low-potential pathway, encoded by *cbcL,* became more important. Changing pH by 0.5 units was not sufficient to further alter reduction phenotypes, suggesting that the freeze drying-aging process lowered the redox potential of ferrihydrite much more than 30 mV of pH change could overcome.

A general trend emerged during these experiments, consistent with autoclaving, freeze drying, and pH influencing redox potential in similar ways, regardless of the mineral. When reduction by Δ*imcH* was near wild-type levels, reduction by Δ*cbcL* was typically at its lowest. When reduction by Δ*cbcL* was near wild-type levels, Δ*imcH* cultures failed to reduce the provided electron acceptor. This trend suggests that 1) the two mutants act as an *in vivo* sensor of initial redox potential, across a suite of minerals, metals, and conditions, and 2) that both systems are employed by wild type bacteria encountering these minerals under normal conditions.

Figure 4 represents all reduction data after seven days of incubation, from 17 different XRD-characterized mineral forms performed in triplicate with three different strains, ranked according to Δ*imcH* performance. Because redox potentials of these minerals can not directly measured, the order shown in Figure 4 cannot be directly related to a quantitative measure of actual redox potential. However, the data provides insight into how simple laboratory manipulations can affect experimental outcomes, and how these are linked to redox potential changes.

**Figure 4.**
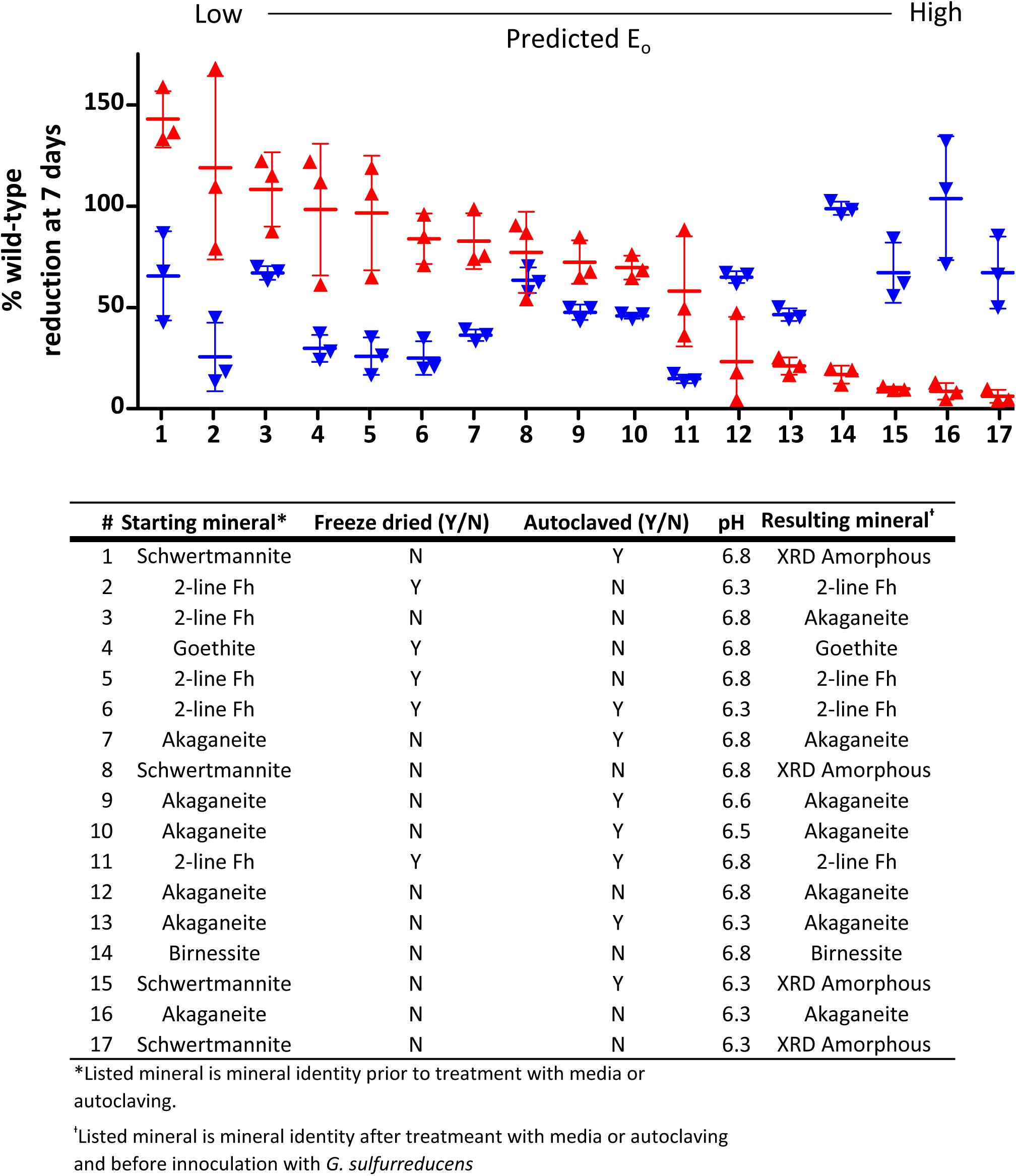
G. sulfurreducens *requires both ImcH and CbcL for complete reduction of a range of minerals*. Cultures of Δ*imcH* (red upward pointing triangles) and Δ*cbcL* (blue downward pointing triangles) were incubated with minerals as the sole terminal electron acceptor, and the extent of Fe(II) or Mn(II) accumulation was measured relative to that of wild type *G. sulfurreducens*. Data shown are the mean and standard deviation of triplicate cultures. Conditions are ordered left to right in order of Δ*imcH* reduction, from best to worst. Akaganeite, β-FeOOH. Schwertmannite, Fe_8_O_8_(OH)_6_(SO_4_)∙nH_2_O. 2-line Fh (Ferrihydrite), ca. Fe_10_O_14_(OH)_2_. Birnessite Na_x_Mn_2-x_(IV)Mn(III)_x_O_4_, x ~ 0.4, Goethite, α-FeOOH.

For example, when Δ*imcH* mutants lacking the high potential pathway were unable to respire, initial conditions were likely well above 0 V vs. SHE, such as with freshly precipitated minerals in medium at pH 6.3. When exposed to the same high potential minerals, Δ*cbcL* mutants containing only the high potential pathway were able to perform similar to wild type. As minerals were aged or manipulated to lower their initial redox potential, the CbcL-dependent pathway became increasingly important to achieve wild-type extents of reduction. These incubations highlighted the different contributions of ImcH and CbcL to the initial rate vs. final extent of Fe(III) reduction, respectively.

Interestingly, some level of metal reduction was always observed for Δ*cbcL* mutants exposed to minerals predicted to be very low potential acceptors. This result is similar to previous studies where Δ*cbcL* still demonstrated a slow but detectable rate of respiration on electrodes poised at low redox potentials (Zacharoff et al., 2016). This activity may reflect a yet-undiscovered pathway used under thermodynamically challenging conditions, where neither ImcH- nor CbcL-dependent pathways can function.

The results in Figure 4 represent a suite of well-understood minerals (2-line ferrihydrite, goethite, akaganeite, schwertmannite, and birnessite), in which variability is typically introduced through freeze-drying, autoclaving, and pH adjustments. These variables alter mineral crystal structure, particle size, degree of aggregation, surface area, and surface charge, and thus influence microbial attachment or behavior in unpredictable ways (Cutting, et al., 2009; Majzlan et al., 2004; Navrotsky et al., 2008; Roden, 2006). However, these manipulations change redox potential in predictable ways. Based on the overall pattern of responses, we propose that redox potential is a master variable affecting the pathway of extracellular electron acceptor reduction used by *G. sulfurreducens.*

### Cells lacking the CbcL-dependent pathway have increased growth yield

A key question stemming from this work relates to the evolutionary forces selecting for multiple electron transfer pathways at the inner membrane. As cells lacking the ImcH-dependent pathway are often able to completely reduce a provided mineral after a small amount of Fe(II) accumulates, why not encode a single electron transfer pathway and allow it operate at all potentials, thus eliminating the need to encode pathways controlling respiration across multiple redox potential windows? One hypothesis is that the multiple pathways encoded *G. sulfurreducens* are able to take advantage of the different amounts of energy represented by the spectrum of metal (oxyhydr)oxides, by more efficiently conserving these energy differences.

To test this hypothesis, the cell yield of *G. sulfurreducens* strains in which one pathway was removed was measured. Because Δ*cbcL* can initially reduce akaganeite, direct comparisons were possible when a low inoculum of each strain was added to medium containing this mineral (autoclaved at pH 6.8, as in Figure 2B) and incubated. In all experiments, Δ*cbcL* generated more colony forming units per mol Fe(II) produced than wild type *G. sulfurreducens.* For each mM of Fe(III) reduced, Δ*cbcL* produced 2.5 ± 0.3 × 10^6^ CFU, while wild type produced 0.8 ± 0.1 and 1.6 ± 0.1 × 10^6^ CFU, respectively (n=5 for each strain) (Figure 5A). These results indicated that when electron flow was forced through the ImcH-dependent pathway (as in Δ*cbcL*), more cells were produced per Fe(III) reduced.

**Figure 5.**
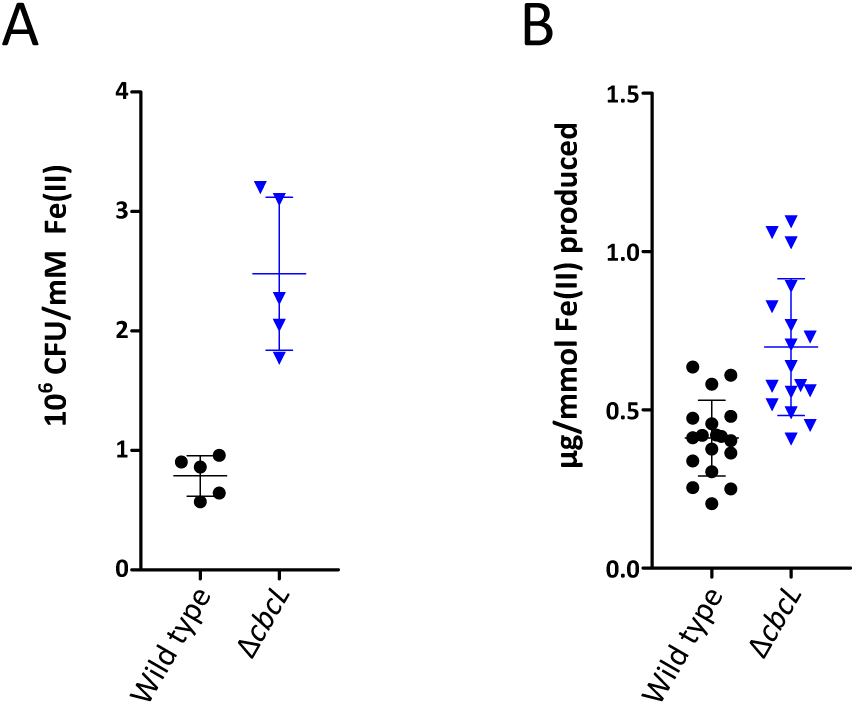
*Δ*cbcL *has an increased growth yield when respiring extracellular electron acceptors*. Removal of the CbcL588 dependent pathway increases the growth yield compared to wild type for two different electron acceptors. (A) For wild-type and Δ*cbcL* strains reducing akaganeite (Autoclaved at pH 6.8, as in Figure 3B) as the sole terminal electron acceptor, Δ*cbcL* generated more cells per unit Fe(II) produced than did wild type. (B) Δ*cbcL* produced more protein per unit Fe(II) produced than wild type when reducing Fe(III)-citrate as the sole terminal electron acceptor. These data suggest that electron flow through ImcH supports conservation of more energy, when high potential electron acceptors reducible by this system are available. Each data point (Black circles=wild type, blue triangles=Δ*cbcL*) represents an individual culture and bars are the mean and standard deviation of the data shown.

Cell attachment to mineral particles could confound plate counts in experiments with cells grown on solid particles. In a second series of experiments, medium containing acetate and fumarate was inoculated with cultures at the same stage of akaganeite reduction. Because all wild-type and mutant cultures grow at similar rates with fumarate as the electron acceptor (see Table 1), the point at which optical density is detectable is a function of the number of cells in the inoculum. In all of these experiments, Δ*cbcL* again demonstrated significantly more cells per mol Fe(II) reduced than wild type.

Yield was also measured after a small inoculum of wild type or Δ*cbcL* cells were allowed to reduce Fe(III)-citrate, and cells were harvested and washed free of soluble iron (which can affect the BCA assay). The Δ*cbcL* cultures again had a significantly increased protein yield (Figure 5B). When standardized to the amount of Fe(II), the yield of cells forced to use only the ImcH pathway was 170% of the wild type using both pathways during complete reduction of Fe(III).

Combined with previous data showing that cells using ImcH-dependent pathways support faster growth rates (Zacharoff et al., 2016), this supports the hypothesis that *G. sulfurreducens* alters its electron transfer pathway to obtain a higher yield per electron transferred. These data indicate that these electron transfer pathways are able to switch ‘on’ and ‘off’ in response to changing redox potential of external electron acceptors. This provides one possible explanation for the complexity and burden associated with encoding multiple pathways of electron transfer across the inner membrane.

## Discussion

Fe(III)-(oxyhydr)oxide minerals in nature exist as a complex continuum of potential energies. As such, it is unsurprising that bacteria would evolve equally complex mechanisms able to harness the energy available during respiration to these metals. While a single pathway essential for reduction of all metals emerged in the model metal-reducing *Shewanella spp.,* a similarly simple solution remained elusive in *Geobacter spp.*. The data reported here demonstrates that even under the most common laboratory conditions where Fe(III)-(oxyhydr)oxide is precipitated and added to medium, *G. sulfurreducens* will utilize multiple electron transfer pathways to accomplish what is considered to be wild-type levels of Fe(III) reduction. For Fe(III)-(oxyhydr)oxides that are predicted to have redox potentials around 0 V, such as schwertmannite, akaganeite, and ferrihydrite, the electron-accepting potential decreases as Fe(II) accumulates, and triggers utilization of electron transfer pathways that support lower cell yield and slower growth rate, but still allow some respiration. The ability of *G. sulfurreducens* to respond to the changing redox potential of its environment likely allows this organism to make the most efficient use of the provided mineral.

Based on proteomic studies, *G. sulfurreducens* expresses at least 78 multiheme *c-*type cytochromes under different laboratory conditions, yet mutant analyses were initially unable to link particular cytochromes to reduction of specific extracellular electron acceptors (Aklujkar et al., 2013; Leang et al., 2003, 2010; Rollefson et al., 2009, 2011; Shi et al., 2007). One hypothesis for cytochrome diversity that drives such -omic comparisons is that separate pathways will exist based on solubility (chelated vs. oxide) or metal (Fe vs. Mn). However, despite their many physiochemical differences, Fe(III)-citrate and Mn(IV)-(oxyhydr)oxides both represent high potential acceptors, from the viewpoint of the inner membrane, and our data suggest that this variable influences at least one portion of the electron transfer pathway. As we detect evidence for at least one other pathway in this work, and the *G. sulfurreducens* genome appears to encode at least 4 other inner membrane quinone oxidoreductases, redox potential differences may help to explain additional complexity of *Geobacter.* Possessing protein machinery able extract the full advantage from minerals as they descend through the redox tower also helps explain the dominance of these metal-reducing organisms in contaminated subsurface environments undergoing rapid redox potential changes.

In order to characterize and biochemically dissect extracellular electron transfer in dissimilatory metal reducing organisms such as *G. sulfurreducens,* closer attention may need to be paid not just to crystal structure, but the actual redox potential experienced by the organisms. Commonly synthesized Fe-(oxyhydr)oxides prepared by precipitation of Fe(III) can produce very different rates of reduction or mutant phenotypes depending on the age of the material and the length of time one is willing to incubate cells. A distinction between initial rates of Fe(III)-(oxyhydr)oxide reduction, where higher-potential conditions exist, and final extent of reduction achieved by slower but lower potential pathways may help separate these confounding effects. As Fe(II) begins to accumulate during the course of Fe(III)- (oxyhydr)oxide reduction by metal reducing microorganisms, the redox potential will change as the ratio of Fe(II)/Fe(III) changes and as mineral structure is altered in response in increased Fe(II) concentration. One would also expect significant differences in washed cell suspensions, where the degree of respiratory coupling to growth is not part of the assay, since electron flow through the ImcH- and CbcL-dependent pathways support different growth rates and yields.

It now appears that in *G. sulfurreducens* a transition from an ImcH-dependent electron transfer pathway to a CbcL-dependent pathway occurs both when electrodes and commonly used Fe(III)-(oxyhydr)oxides are provided as sole terminal electron acceptors. The initial rate and total extent of Fe(III)-(oxyhydr)oxide reduction may also be the result of a series of redox potential dependent processes in environmental samples, with each step supporting different cell yields. Competition in the environment could occur along any part of this continuum, rewarding those capable of rapid growth when the Fe(III)-(oxyhydr)oxide is high potential, or those able to survive with electron acceptors at relatively low redox potentials. The prevalence of ImcH and CbcL homologues in other dissimilatory metal reducing organisms and in metagenomics data from sites undergoing active metal reduction (Levar et al., 2014; Zacharoff et al., 2016) suggests that this type of redox discrimination could also occur in diverse organisms, and provides opportunities for further exploration of redox potential dependent respiration in these organisms and their environments.

## Acknowledgements

C.E.L. was supported by the State of Minnesota’s MNDrive program, D.R.B. was partially supported by the Office of Naval Research grant N000141210308, A.J.D was supported by UMN U.R.O.P, and C.L.H and B.M.T were supported by the State of Minnesota’s L.C.C.M.R program.

